# Multi-omics time-series analysis in microbiome research: a systematic review

**DOI:** 10.1101/2025.07.03.659054

**Authors:** Moiz Khan Sherwani, Matti O. Ruuskanen, Dylan Feldner-Busztin, Panos Nisantzis Firbas, Gergely Boza, Ágnes Móréh, Tuomas Borman, Pande Putu Erawijantari, István Scheuring, Shyam Gopalakrishnan, Leo Lahti

## Abstract

Recent developments in data generation have opened up unprecedented insights into living systems. It has been recognized that integrating and characterizing temporal variation simultaneously across multiple scales, from specific molecular interactions to entire ecosystems, is crucial for uncovering biological mechanisms and understanding the emergence of complex phenotypes. With the increasing number of studies incorporating multi-omics data sampled over time, it has become clear that integrated approaches are pivotal for these efforts. However, standard data analytical practices in longitudinal multi-omics are still shaping up and many of the available methods have not yet been widely evaluated and adopted. Thus, we performed a systematic literature review on data science methods for longitudinal multi-omics, with a particular focus on microbiome research.

## Introduction

Multicellular organisms coexist with microbes, collectively constituting a *holobiont* (1). For a holistic understanding of the host organism, we need to understand the network of interactions between the host and its microbiomes. Some of the most important questions related to multicellular hosts are e.g., how they maintain their homeostasis, react to changing environments, defend against infections and how the interactions between the hosts and their microbiomes contribute to these processes. To answer these questions, we should analyse the host genome (complete DNA sequence), epigenome (chemical modifications to DNA), transcriptome (all RNA transcripts), proteome (set of proteins expressed in an organism), metabolome (complete set of metabolites) and other aspects of the system in parallel. Thus, the collection of multi-omics data from the host and host-associated microbiomes and the development of multiomics analysis techniques has emerged as an active research topic (2). Despite the progress, revealing causal relations and accounting for temporal variation in multi-omics studies necessitates sampling across different time points and treatment conditions.

Multi-omics data and the related analysis methods are heterogeneous. The various ’omics represent very different types of biological molecules. Metagenomics involves the comprehensive sequencing of all microbial genomes within a sample, enabling the reconstruction of functional potential and the community structure of the microbiome (3). Transcriptomics measures RNA transcripts to estimate the relative expression of genes (4). Proteomics includes quantitative measures of the different proteins (5), while metataxonomics aims to characterise all microbial taxa in a sample (6), often through 16S rRNA gene amplicon sequencing (7). Genomics enables us to determine whether mutations are present at specific positions in the genome and epigenomics informs us on differences in gene regulation (8). Each of these methods has its own technical challenges and resulting biases, for example, the identification and quantification of proteins with low abundance in proteomics (5), selecting an appropriate pre-processing method in transcriptomics (9) and deciding how to define the units of analysis in metataxonomics (10). There are also further issues of high dimensionality (large number of features or variables compared to the relatively small number of samples), high stochasticity or noise (random variations that obscure true signal) and batch effects (systematic variations introduced by differences in experimental conditions). Taken together, these and other properties of the data can make the interpretation and use of the data difficult (11). Furthermore, matching the samples and features between the complementary ’omics is necessary for joint analysis but not always straightforward. A description of these challenges has been given by Chalise et al. (12).

Including the temporal dimension brings in an additional layer of challenges in terms of data collection and analysis. It might be impossible to comprehensively analyse the same entity, such as a developing organism, at different time points. In such cases, it might be necessary to perform a pseudotime series, i.e. a time-series with different samples from a (relatively) homogeneous population sampled at different time points. Additionally, temporal multiomics can provide various benefits, such as balancing out individual variability (13) and provide a dynamic view on the holobiont.

Thus, to comprehend the multitude of interactions occurring within the holobiont over time, it is necessary to employ temporal version of multi-omics analysis techniques. This review provides a systematic overview of data analytical methods in longitudinal multi-omics and highlights emerging topics for future research.

## Systematic review method

### Preliminary systematic search and screening of studies

We selected the reviewed studies based on the PRISMA guidelines (14) (Fig. 1) and with specific inclusion/exclusion criteria (Fig. 2). We defined multi-omics data as a combination of two or more ’omics datasets that included longitudinal measurements as real or pseudo time-series. It was vital to include research that include pseudo-time series since sampling can sometimes be destructive, which makes it hard to collect full longitudinal observations (15). The search for relevant literature was conducted on June 15th, 2024, using the key “multi-omics (“time series” OR “over time” OR “temporal” OR “longitudinal”)” in all domains of the Web of Science. The Scopus database was queried using the search term “TITLE-ABS-KEY(“multi” AND “omics” AND (“time series” OR “across time” OR “temporal” OR “longitudinal”))”. We restricted the searches to original studies written in English and excluded review studies. This yielded 382 entries from Web of Science and 459 entries from Scopus. After manually identifying and eliminating 311 duplicates, 530 distinct records remained for analysis. Based on abstract screening, we excluded studies that did not align within the defined scope. The remaining 174 studies underwent full-text screening, during which we excluded further 31 studies out of 174 studies due to incomplete information as defined in our study design Fig. 2. A total of 143 studies fulfilled the criteria established for this review.

**Fig. 1.**
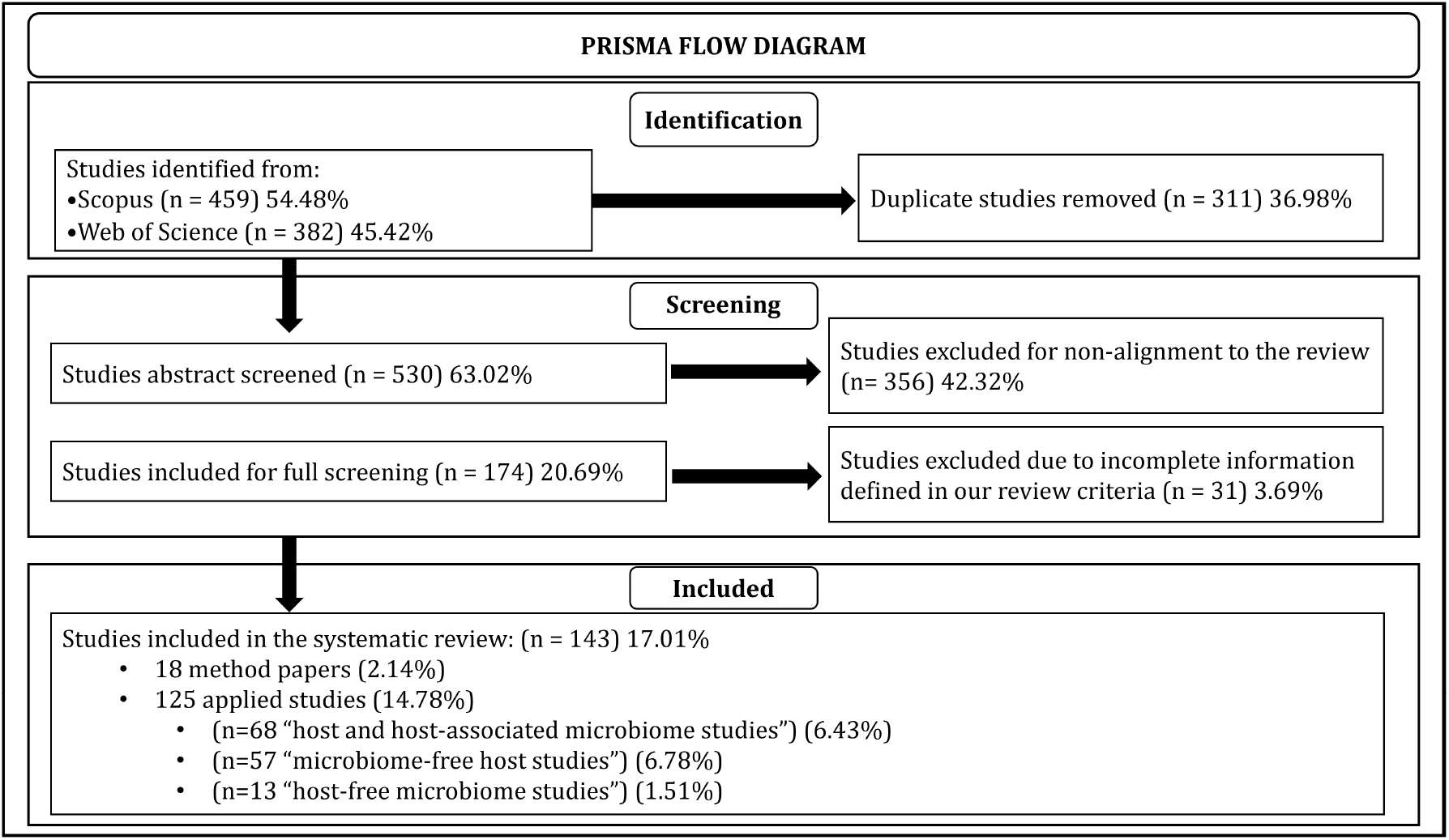
PRISMA flow diagram illustrating the study selection process for this review.

**Fig. 2.**
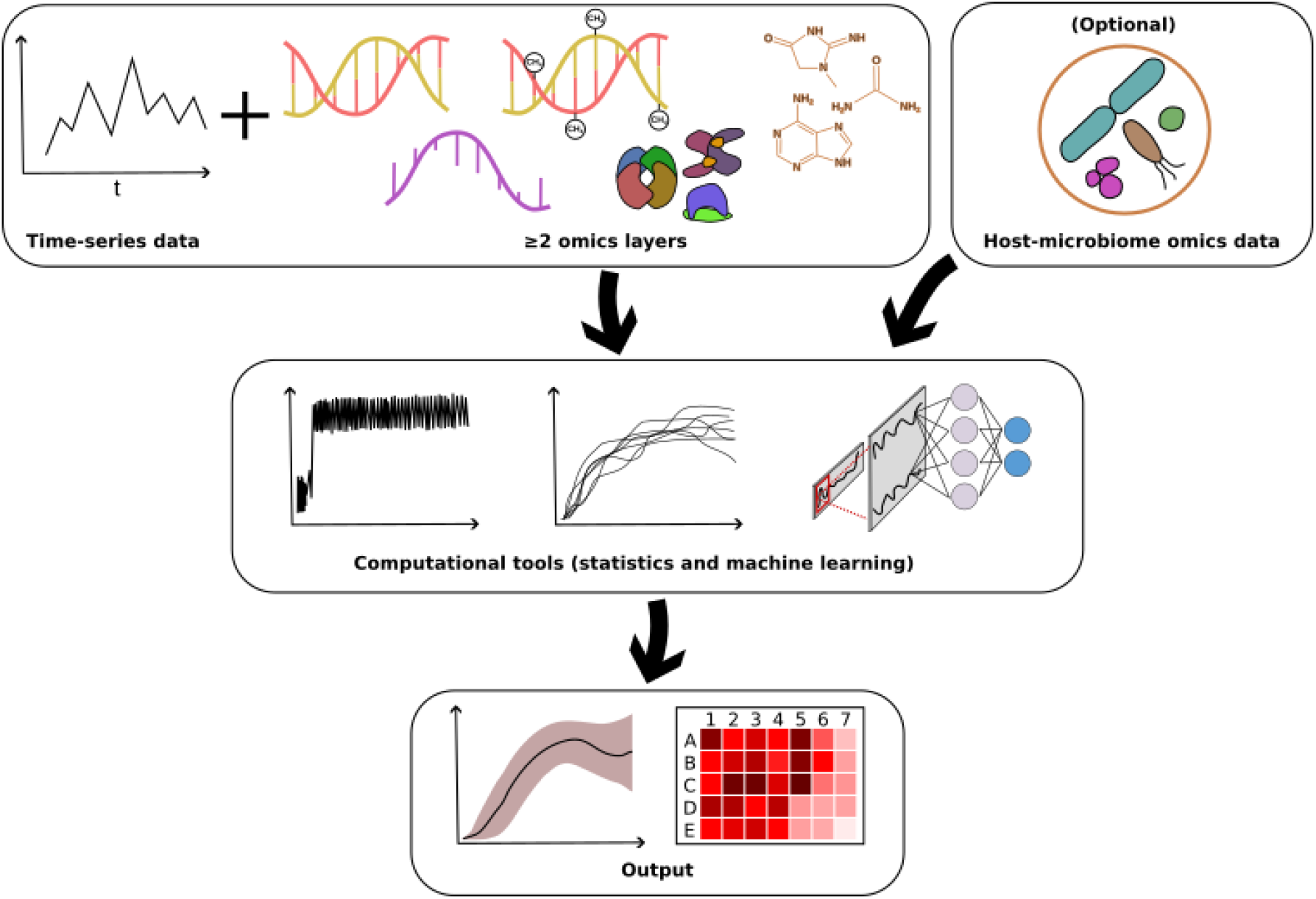
Overview of the study design in this systematic review. The architecture illustrates the integration of multi-omics time-series data layers (min. 2 layers), host-associated microbiome data (optional), into computational analysis using statistical, machine learning (ML) and deep learning (DL) methods to obtain the output. The models can output e.g., various types of predictions and correlations. The diagram outlines the process flow from data collection to computational processing and final output generation, showing the methodological framework used in the reviewed studies.

### Evaluation of studies

The 143 studies that met our systematic review criteria consisted of 125 (87%) applied studies and 18 (13%) methodological studies. Among the 125 applied studies, 55 included “host and host-associated microbiome data” - these studies investigate both the host and its associated microbial communities (Table 1). Of the remaining 70 studies, 57 included “microbiome-free host data” - these studies focus exclusively on the host, analyzing host genomics, transcriptomics, proteomics, or metabolomics without considering microbial data (Table 2). Finally, there were 13 studies that focused on “host-free microbiome data” - these studies examine microbial communities in environments or contexts where a host is not involved, such as free-living or environmental microbiomes (Table 3). Each study was evaluated by at least two authors to ensure consistency. For the applied studies, we systematically summarized key aspects, including the types of samples analysed, the frequency and duration of sampling, the types of ’omics data used and the analytical approaches employed. For the methods-based studies, we performed a qualitative assessment based on established criteria: predictive performance, interpretability and ease of installation/use. These criteria were selected to address fundamental considerations for the actual implementation and usability of methods in multi-omics research. Predictive performance was emphasized to guarantee the reliability and accuracy of results, while interpretability evaluated the capacity of each approach to produce understandable and significant insights for users. User-friendly implementation and the ease of installation are essential considerations for the accessibility of a wider scientific audience. Furthermore, we evaluated the development and maintenance activities of each technique in order to further assess the robustness and long-term availability. This assessment included data from the primary studies and associated resources, including code repositories, tutorials and online documentation (Table 4). Collectively, these factors underscore the necessity of choosing methods that are robust, pragmatic and consistently maintained.

**Table 1.**
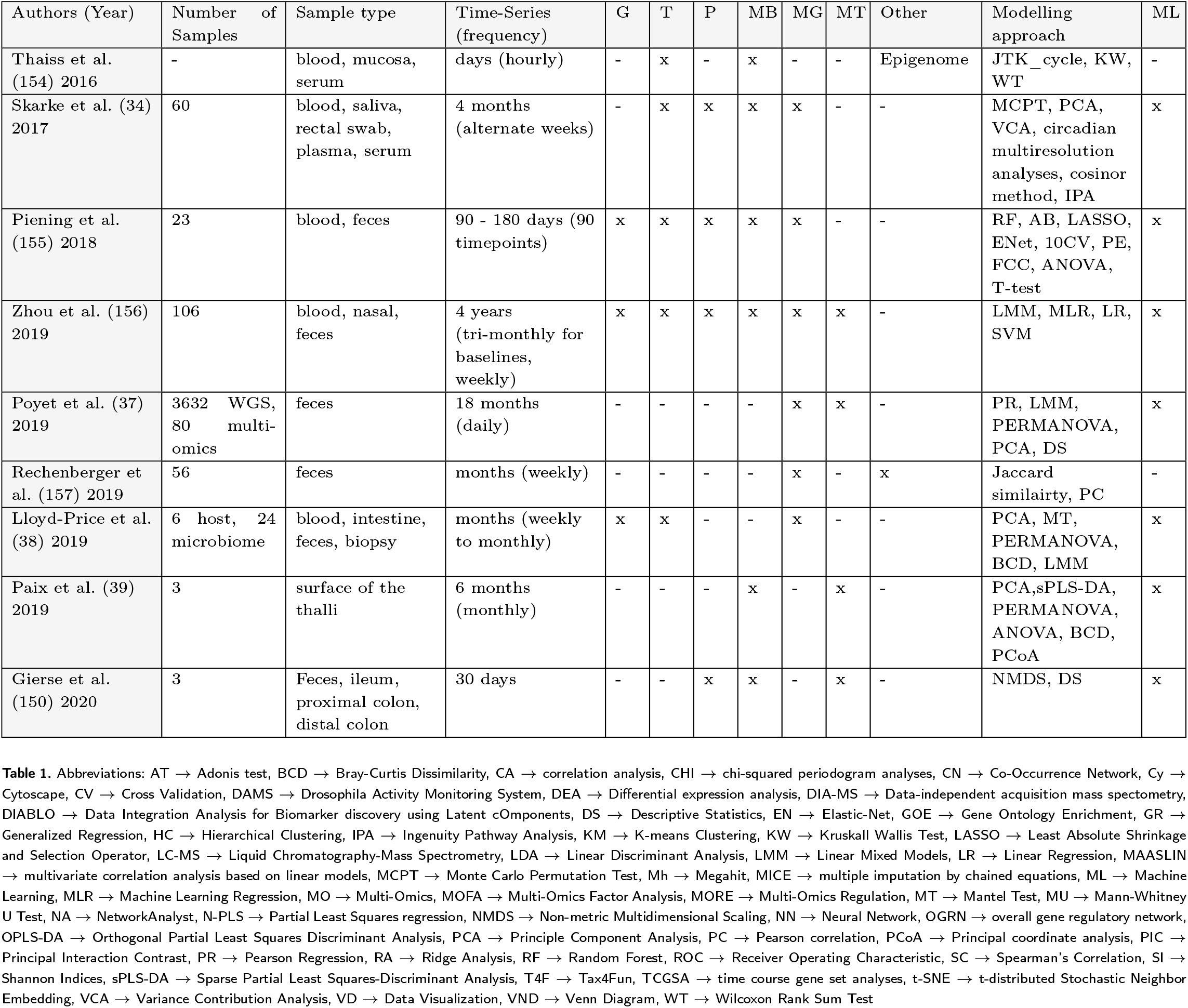
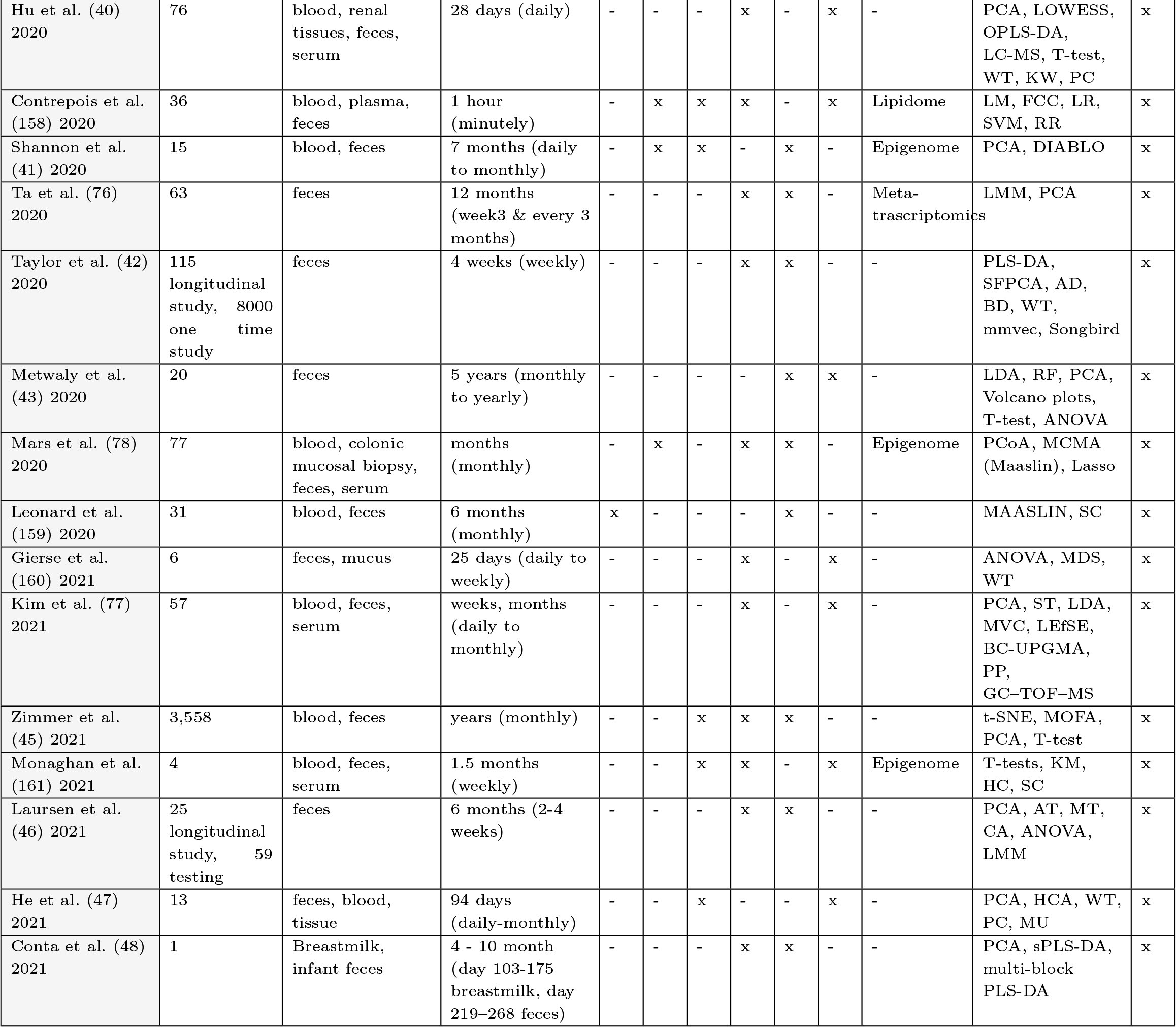
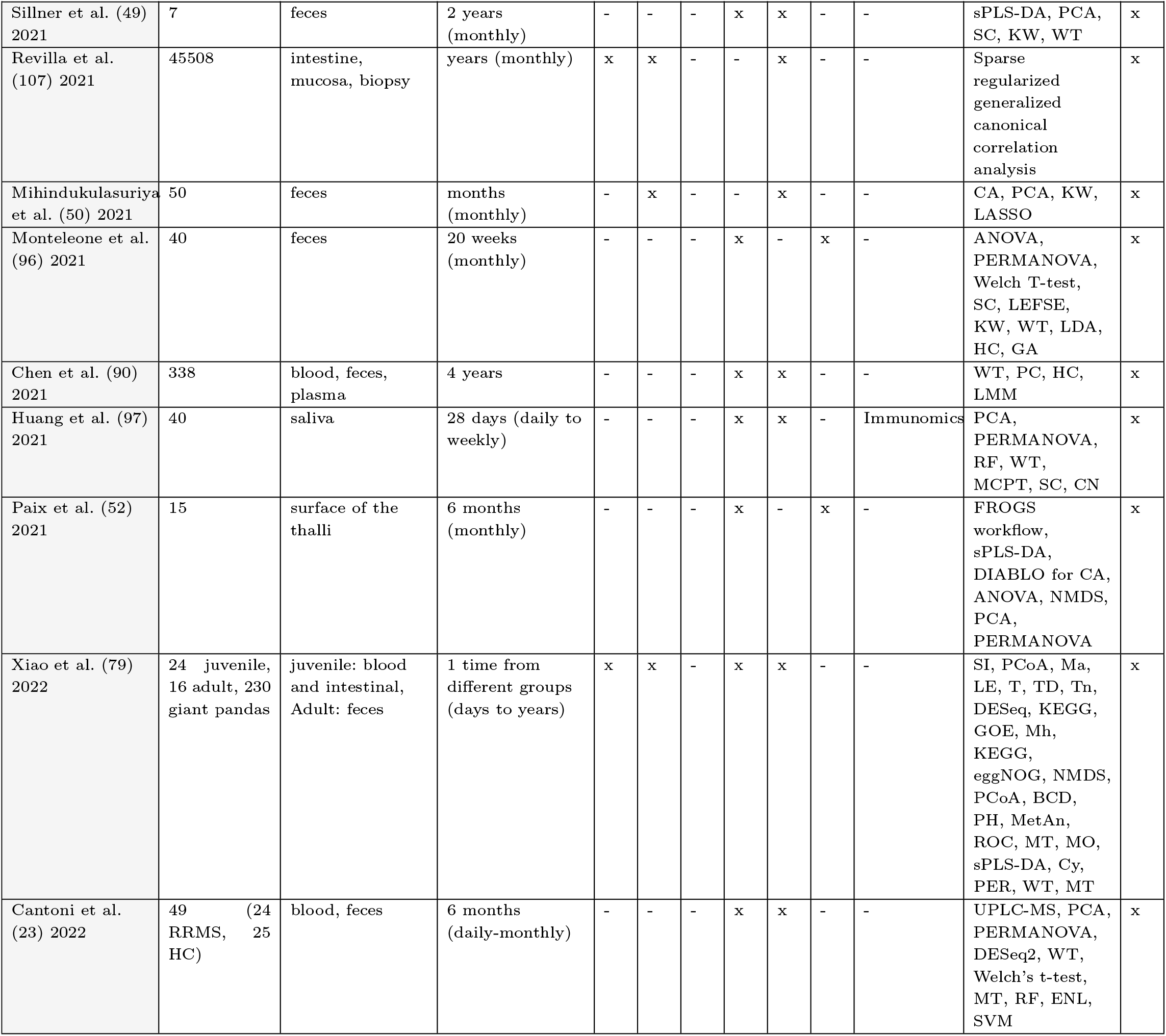
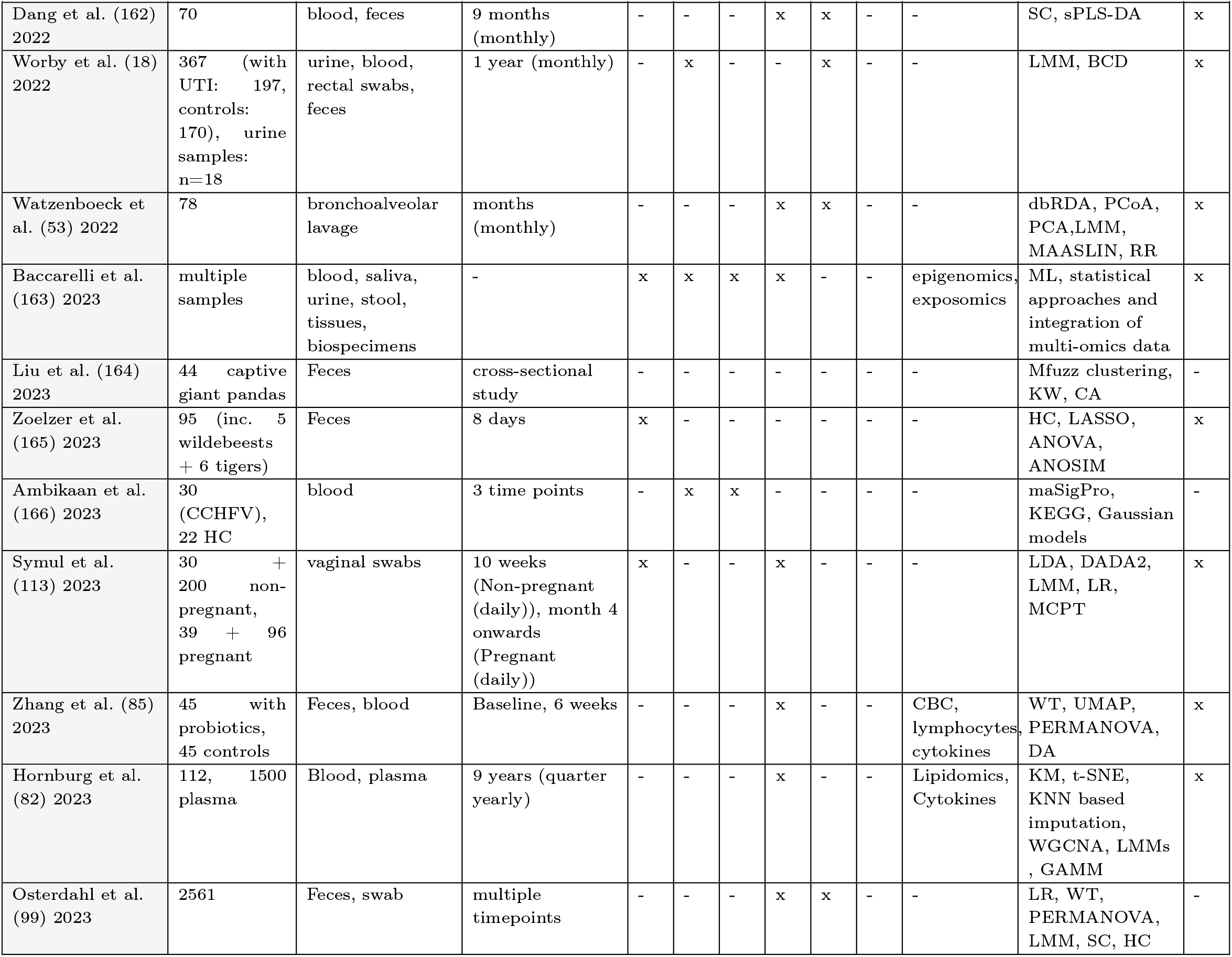
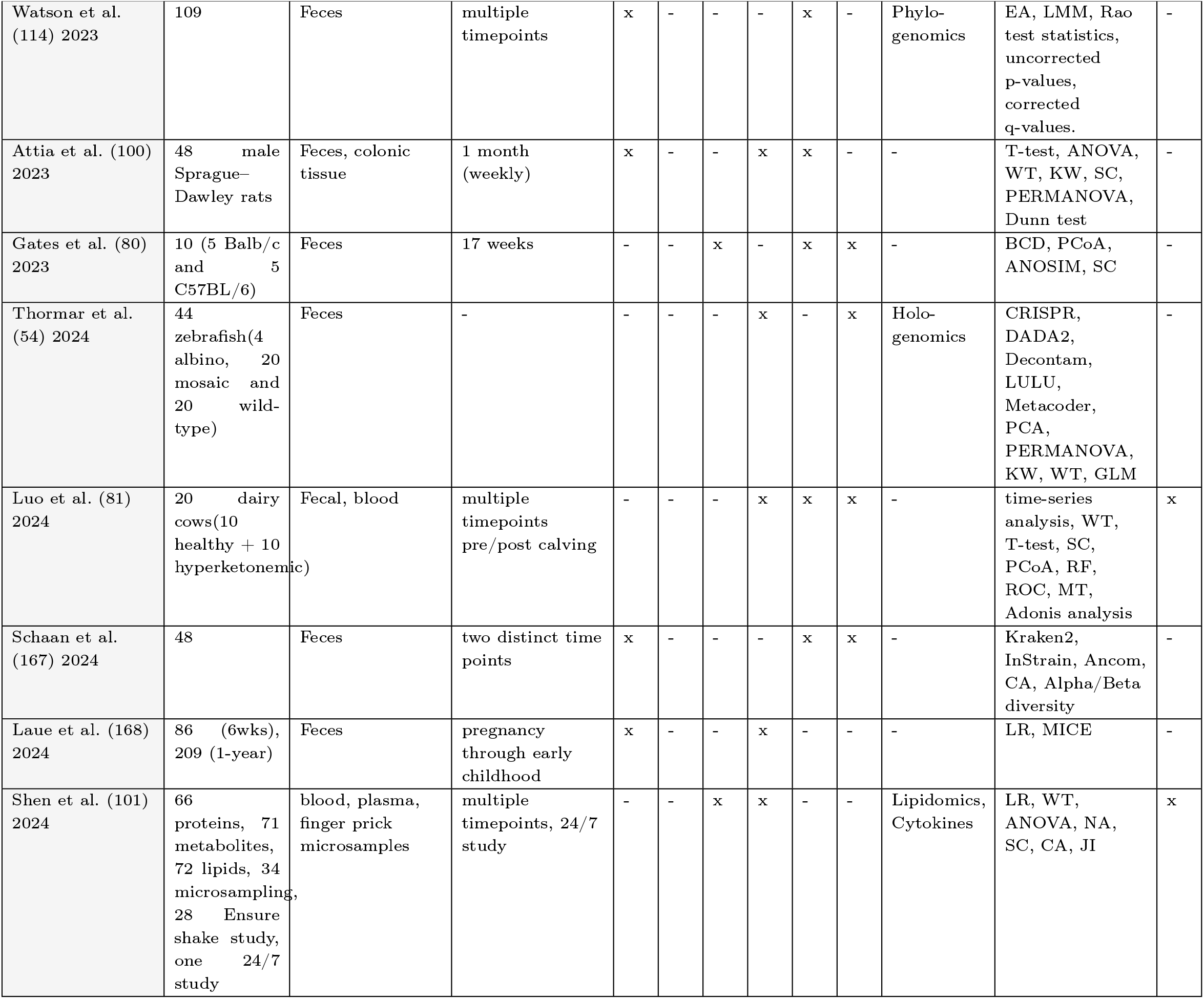
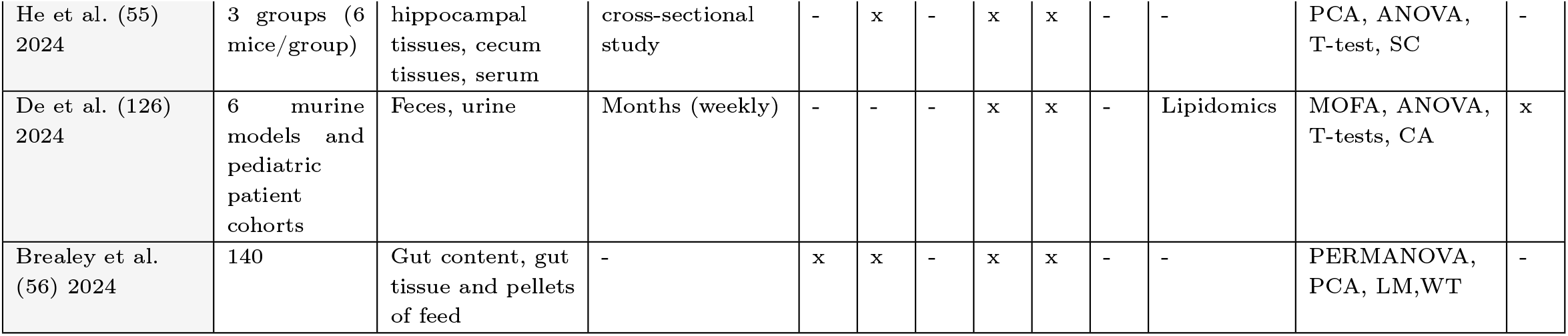
Overview of multi-omics “host and host-associated microbiome” studies: including authors, year of study, sample type, temporal sampling frequency, data types (Genomic, Transcriptomic, Proteomic, Metabolomic, Metagenomics, Metataxonomics and Others) and applied modeling or machine learning approaches.

**Table 2.** Overview of multi-omics “microbiome-free host” studies: including authors, year of study, sample type, temporal sampling frequency, data types (Genomic, Transcriptomic, Proteomic, Metabolomic and Others) and applied modeling or machine learning approaches.

| Authors (Year) | Number of Samples | sample types | Time-Series (frequency) | G | T | P | MB | Others | Modelling approach | ML |
| --- | --- | --- | --- | --- | --- | --- | --- | --- | --- | --- |
| Ansong et al. (169) 2013 | 3 | cell cultures | 8 hours (hourly) | - | x | x | x | Metagenome | microarray analysis, LC-MS, NMR, GC-MS, context likelihood of relatedness, Louvain-community-finding algorithm | x |
| Kihara et al. (170) 2014 | 15 | cell cultures | (hourly) | - | x | - | - | Lipidome | linear kinetics, ODE model | - |
| Gong et al. (171) 2015 | 4 | cell cultures | 4 timepoints | - | x | - | - | Epigenome | LR, Bayesian network model | x |
| Tan et al. (172) 2017 | 8 | cell cultures | 16 hours (hourly) | - | - | x | - | Phospho-proteome | ANOVA, WCGNA for NA | x |
| Harvald et al. (91) 2017 | 42 | whole organism | 16 hours (hourly) | - | x | x | - | - | LC-MS, PC, HC, KEGG, GOE | x |
| Shih et al. (21) 2017 | 1205 AN + 1948 control | blood | - | x | - | x | x | Lipidome | LC-MS, normality tests, ANOVA | - |
| Ahn et al. (173) 2017 | 27 | whole plants | 6 hours (multiple timepoints) | - | x | - | - | Epigenome | TF networks, Cascade tree, T-test, Fisher's test | - |
| Sánchez-Gaya et al. (174) 2018 | 16 | cell cultures | - | - | x | - | - | Epigenome | N-PLS, MORE | x |
| Tasaki et al. (19) 2018 | hundreds | blood, cell cultures | years (weekly to monthly) | - | x | x | - | - | PLSR | - |
| Sarigiannis et al. (175) 2018 | 350 children | urine | multiple timepoints | x | x | x | x | Epigenome | Correlation globe plots for associations using effect size | - |

**Table 2.** List of Abbreviations: AT → Adonis test, BCD → Bray-Curtis Dissimilarity, CA → correlation analysis, CHI → chi-squared periodogram analyses, CN → Co-Occurrence Network, Cy → Cytoscape, CV → Cross Validation, DAMS → Drosophila Activity Monitoring System, DEA → Differential expression ...
| Authors (Year) | Number of Samples | sample types | Time-Series (frequency) | G | T | P | MB | Others | Modelling approach | ML |
| --- | --- | --- | --- | --- | --- | --- | --- | --- | --- | --- |
| Abreu et al. (57) 2018 | 171 seedlings, 157 leaf samples | seedlings, leaves | 12 hours (multiple timepoints) | - | x | - | - | Lipidome | graph-guided fused least absolute shrinkage and selection operator, PCA, PC, NA | x |
| Sumit et al. (58) 2019 | 4 | cell cultures | days (daily) | - | x | - | x | - | PCA, GSEA, TCGSA, maSigPro | x |
| Pavkovic et al. (59) 2019 | FA model: 5x2; UUO: 4x4(day 0: 3) | kidney tissue | 2 weeks (daily) | - | x | x | - | - | FastQC, STAR/Seqbuster, DESEQ, LDA, PCA | x |
| Simats et al. (60) 2020 | 37, 9 excluded | brain tissue | only once | - | x | x | - | - | PCA, Multiple CIA, regularized Canonical CA | x |
| Lin et al. (61) 2020 | 6 | blood, kidney tissue | days (daily) | - | - | x | - | Phospho-proteome | HC, PCA, T-test, ANOVA, NA | x |
| Zhao et al. (176) 2020 | 23 | urine | 18 hours (hourly) | - | - | x | x | - | The analysis were performed individually on each 'omics | x |
| Bernardes et al. (62) 2020 | 14 | blood | weeks (daily) | - | x | - | - | Epigenome | UMAP, PCA, HC | x |
| Wang et al. (177) 2020 | 32 | whole heads | 2 days (Every 3 hours) | - | x | x | - | - | DAMS, two-sided hypergeometric test, CHI | x |
| Seifert et al. (92) 2020 | 22 | tumor tissue | - | x | x | - | - | - | PC, HC, heatmaps, VND, differential expression analysis Fisher's test | - |
| Zander et al. (178) 2020 | tissue samples from more entities | seedlings | hours (minute to hourly) | - | x | x | - | - | GC, RTP-Star package for gene regulatory network | x |
| Lam et al. (179) 2021 | 78 | blood | - | - | x | x | - | - | Pairwise statistical analysis, 'omics was analysed individually | x |
| Tarca et al. (180) 2021 | 133 | blood | between 4-7 weeks (2 timepoints) | - | x | x | - | - | LASSO, RF, RR, GR, SVM | x |
| Suvarna et al. (181) 2021 | 2 | blood | weeks (weekly) | - | - | x | x | Lipidome | IPA | x |
| Yang et al. (182) 2021 | - | blood | - | - | - | - | - | - | - | x |
| Brands et al. (20) 2021 | 76 (56 after 1 month), 41 CP | blood | 1 month | - | x | - | - | Epigenome | MAASLIN, DIABLO | x |
| Matsuzak et al. (63) 2021 | 9 per time point | liver, blood | 4 hours (minutes) | - | x | x | x | - | DS, PCA, HC, NA | x |
| Sprengr et al. (93) 2021 | 3 per time point | liver tissue, blood | 48 hours (hourly) | - | - | x | - | Lipidome | ANOVA, PC, c-means clustering, HS, T-test | x |
| Wu et al. (16) 2021 | 194 (+472 Validation (trauma dataset 2 [TD2])) | blood | days (daily) | - | - | x | x | Lipidome | HC, KM | x |
| Djeddi et al. (64) 2021 | 36 | blood | 7 weeks (weekly) | - | x | x | - | - | PCA, T-test | x |
| Lee et al. (94) 2021 | 3 bioreactors | cell cultures | 14 days (daily) | - | x | x | x | - | T-test, ANOVA, enrichment analysis, Fisher's test, HC, PC, heatmap | x |
| Schwaber et al. (183) 2021 | 1 (triplicate samples from one culture) | cell cultures | 9 days (daily) | - | x | x | - | - | HC, NA, heatmap | x |
| Liu et al. (83) 2021 | 60 | blood | days (daily) | - | x | x | x | x | HC, N-PLS, UMAP, ANOVA | x |
| Balzano-Nogueira et al. (184) 2021 | 306 | blood | 12 months (monthly) | x | x | - | x | Epigenome | NPLS-DA, CV, Partial correlation, GSEA, multi-omics data visualization | x |
| Sun et al. (65) 2021 | 33 | blood | weeks (daily) | - | x | x | x | Lipidome | PCA, functional enrichment analysis, KM, CN, heat map | x |
| Tang et al. (116) 2021 | 76 | blood, urine, fingernails | days (daily) | - | x | - | x | Epigenome, Lipidome | DS, sPLS-DA, RF, Multivariate Analysis, CIT, GLM, ROC | x |
| Liu et al. (83) 2021 | 14 patients, 12000 plasma, 57000 immune cells | blood, cell cultures | 2-4 timepoints | x | x | - | - | - | t-SNE, gene expression profiles at different time points | x |
| Codrich et al. (185) 2021 | - | cell cultures | (hourly) | x | x | x | - | - | Mutect2, HC | x |
| Rodrigues et al. (66) 2021 | - | cell cultures | 72h (hourly) | x | x | - | x | - | COSMOS using prior knowledge, PCA, NA, time-dependent gene clustering, flow injection-MS | x |
| Singhal et al. (67) 2021 | 4 | lung endothelial cells, blood | 36 days (weekly) | - | x | x | - | - | PCA, heatmap, MU | x |
| Clark et al. (68) 2021 | 4 seedlings per time point | seedlings | 8 hours (minutes to hourly) | - | x | x | - | - | DEA, HC, PCA, PoissonSeq, NA, PC, SC, GLMs | x |
| Camargo et al. (186) 2021 | 8 | leaves, shoot tissue | 26 days (daily) | - | x | - | - | - | GSEA, DESEQ, KM, LASSO, discrimination of gene network structures | x |
| Sacco et al. (95) 2022 | 186 | blood | 7 days (days) | x | x | x | - | Epigenome | PC, RF | x |
| Zoran et al. (187) 2022 | 6 | blood | 2 months (daily to weekly) | - | x | - | x | - | MU, heatmap, GSEA | - |
| Neogi et al. (69) 2022 | 12 | blood | years | - | x | x | x | - | DESEQ, NA, PCA, T-test | x |
| Song et al. (188) 2022 | 3-8 mouse (total 36 samples) + 36 human heart samples) | heart tissue | days (hourly) | - | x | x | x | - | T-test, ANOVA, MT, WT | - |
| Li et al. (189) 2022 | 4 different hematopoietic cell lines through development | cell cultures | between day 10-14.5 (multiple timepoints) | - | x | - | - | Epigenome | TAD, gene expression and correlation with TF binding motif | x |
| Unterman et al. (86) 2022 | 18 pbmc samples + 10 | cell cultures | pre/post covid (weekly to monthly) | - | x | x | - | - | Louvain clustering, UMAP, IgPhyML (lineage tree analysis), WT, MU | x |
| Su et al. (22) 2022 | 209 patients + 457 controls | blood | months (weekly to monthly) | - | x | x | x | - | UMAP, IPA, heatmap, CA, MU, PCA | x |
| Pekayvaz et al. (87) 2022 | 82 | blood, nasal swab | weeks (daily) | x | x | x | - | - | UMAP, Tempora analysis, NA | x |
| Morilla et al. (84) 2022 | 5 | cell cultures | days (daily) | - | x | x | x | - | HC, PC, t-SNE, NN, OGRN, T-test, ANOVA | x |
| Cui et al. (70) 2022 | 300 | whole plants, seedlings | 10 days (twice days apart) | - | x | x | - | Epigenome | DIA-MS, GOE, ANOVA, PCA, box plots, heatmaps, violin plots | x |
| Reimer et al. (71) 2022 | 96 | leaves | 14 days (weekly) | - | x | - | x | - | Weighted cluster analysis, DIABLO, ANOVA, PCA | x |
| Zhang et al. (190) 2022 | tissue samples from more entities | stems, leaves, roots, buds | weeks (days) | x | x | - | x | - | heat maps, gene expressions vs flower stages | x |
| Allesoe et al. (191) 2023 | 789 (T2 diabetes) | blood | (0, 18 & 36) month | x | x | x | - | - | T-tests, ANOVA, VAE, MOVE | x |
| Zheng et al. (192) 2023 | - | Tea leaves inoculated with Pseudopestalotiopsis theae | (0, 1, 3 & 6) day | - | x | - | x | - | Fisher's test, GEA, multivariate testing | - |
| Wang et al. (102) 2024 | Multiple samples | large intestinal tissues from M. fascicularis, cell lines (Caco-2 and HEK293T) and C. elegans were used for cell culture and RNAi | cross-sectional study | - | - | x | x | - | ANOVA, T-tests, KW, SC, PC | - |
| Ciurli et al. (72) 2024 | 10 Male, 10 Female 18 - 45 age | Saliva(above Tongue, below Tongue, right cheek) | 3x/day | - | - | x | x | - | HC, PCA, SC, WT, sPLS-DA | x |

**Table 3.** Overview of multi-omics “host-free microbiome” studies: including authors, year of study, sample type, temporal sampling frequency, data types (Genomic, Transcriptomic, Proteomic, Metabolomic and Others) and applied modeling or machine learning approaches.

| Authors (Year) | Number of Samples | Sample type | Time-Series (frequency) | G | T | P | MB | MG | MT | Other | Modelling approach | ML |
| --- | --- | --- | --- | --- | --- | --- | --- | --- | --- | --- | --- | --- |
| Muller et al. (193) 2014 | 1 | wastewater treatment anoxic phase | 1 year (monthly) | - | x | x | x | x | - | Meta-proteomics, Meta-transcriptomics | WT | x |
| Mannan et al. (194) 2015 | - | bioreactor | - | x | x | x | x | - | - | - | Kinetic modelling | x |
| Alessi et al. (35) 2018 | 3 | compost, wheat straw | 8 weeks (weekly) | - | - | x | - | - | x | - | PCA, ANOVA, VND, HC, MDS | x |
| Han et al. (36) 2018 | 3 | bioreactor | 12 hours (minutes to hourly) | - | x | x | x | - | - | - | PCA, OPLS-DA | x |
| Watahiki et al. (75) 2019 | 2 | groundwater | 3 months (daily to monthly) | x | - | - | - | x | - | - | PCoA, DESEQ, HC | x |
| Wang et al. (89) 2019 | 1 | bioreactor, partial-nitritation anammox reactor | 6 months (weekly to monthly) | - | - | - | - | x | x | - | PC, HC | x |
| Kim et al. (115) 2020 | 1 | bioreactor | 2 days (minute to daily) | - | x | - | x | - | - | - | HC, PCA, VND, ANOVA, sPLS-DA | x |
| Delogu et al. (44) 2020 | pseudo time-series of 3 flasks per time point | bioreactor | 43h (5 hours) | - | x | - | x | x | - | Meta-proteomics | Protein expression control analysis, PC, PCA, LMM | x |
| Breister et al. (195) 2020 | 6 | bioreactor | 24 weeks (weekly) | x | - | - | - | x | x | - | - | x |
| Kralj et al. (196) 2022 | 1 | bioreactor | - | - | - | x | - | - | - | Lipidomics | T-test volcano plot | x |

**Table 3.** Abbreviations: AT → Adonis test, BCD → Bray-Curtis Dissimilarity, CA → correlation analysis, CHI → chi-squared periodogram analyses, CN → Co-Occurrence Network, Cy → Cytoscape, CV → Cross Validation, DAMS → Drosophila Activity Monitoring System, DEA → Differential expression analysis, ...
| Authors (Year) | Number of Samples | Sample type | Time-Series (frequency) | G | T | P | MB | MG | MT | Other | Modelling approach | ML |
| --- | --- | --- | --- | --- | --- | --- | --- | --- | --- | --- | --- | --- |
| Kleikamp et al. (197) 2023 | 3 wastewater treatment plants | Aerobic granular sludge of 2 mm | - | - | - | x | x | - | - | Lipidomics, Cytokines | KEGG, COG terms, PFAM | - |
| Dong et al. (198) 2024 | - | PHE-contaminated soil | 28 days | - | - | - | x | x | - | - | Heatmap, NA | - |
| Delogu et al. (24) 2024 | 51, 21 | floating biomass | 1.5 years(weekly) | - | x | x | - | x | - | - | LR, CA, Ljung-Box test, Kwiatkowski-Phillips-Schmidt-Shin test | x |

**Table 4.**
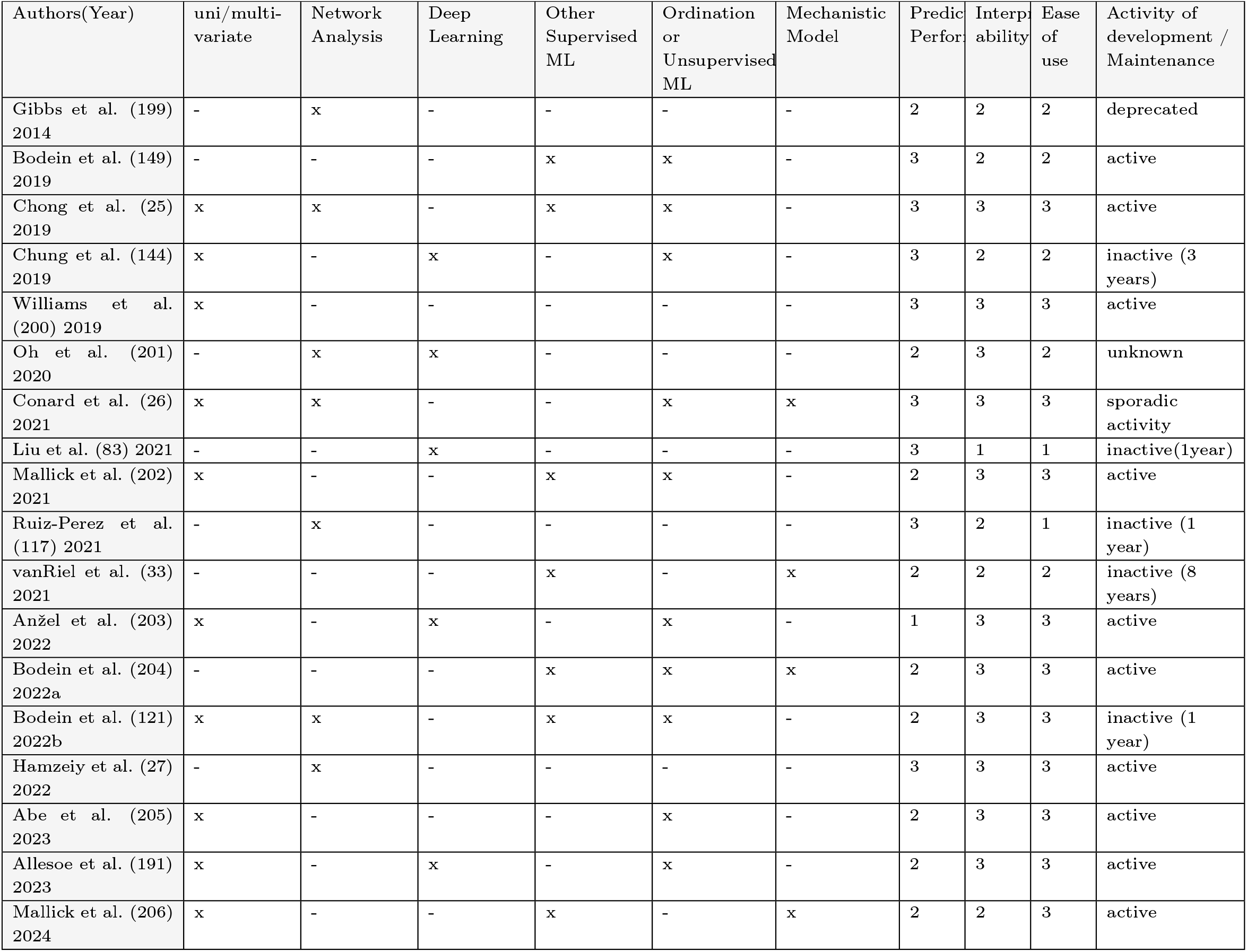
Overview of multi-omics method studies: detailing statistical and machine learning approaches (univariate/multivariate analysis, network analysis, deep learning, supervised and unsupervised ml, mechanistic models), along with predictive performance, interpretability, ease of use and activity status of development or maintenance.

| Authors(Year) | uni/multi-variate | Network Analysis | Deep Learning | Other Supervised ML | Ordination or Unsupervised ML | Mechanistic Model | Predictive Performance | Interpretability | Ease of use | Activity of development / Maintenance |
| --- | --- | --- | --- | --- | --- | --- | --- | --- | --- | --- |
| Gibbs et al. (199) 2014 | - | x | - | - | - | - | 2 | 2 | 2 | deprecated |
| Bodein et al. (149) 2019 | - | - | - | x | x | - | 3 | 2 | 2 | active |
| Chong et al. (25) 2019 | x | x | - | x | x | - | 3 | 3 | 3 | active |
| Chung et al. (144) 2019 | x | - | x | - | x | - | 3 | 2 | 2 | inactive (3 years) |
| Williams et al. (200) 2019 | x | - | - | - | - | - | 3 | 3 | 3 | active |
| Oh et al. (201) 2020 | - | x | x | - | - | - | 2 | 3 | 2 | unknown |
| Conard et al. (26) 2021 | x | x | - | - | x | x | 3 | 3 | 3 | sporadic activity |
| Liu et al. (83) 2021 | - | - | x | - | - | - | 3 | 1 | 1 | inactive(1year) |
| Mallick et al. (202) 2021 | x | - | - | x | x | - | 2 | 3 | 3 | active |
| Ruiz-Perez et al. (117) 2021 | - | x | - | - | - | - | 3 | 2 | 1 | inactive (1 year) |
| vanRiel et al. (33) 2021 | - | - | - | x | - | x | 2 | 2 | 2 | inactive (8 years) |
| Anžel et al. (203) 2022 | x | - | x | - | x | - | 1 | 3 | 3 | active |
| Bodein et al. (204) 2022a | - | - | - | x | x | x | 2 | 3 | 3 | active |
| Bodein et al. (121) 2022b | x | x | - | x | x | - | 2 | 3 | 3 | inactive (1 year) |
| Hamzeiy et al. (27) 2022 | - | x | - | - | - | - | 3 | 3 | 3 | active |
| Abe et al. (205) 2023 | x | - | - | - | x | - | 2 | 3 | 3 | active |
| Allesoe et al. (191) 2023 | x | - | x | - | x | - | 2 | 3 | 3 | active |
| Mallick et al. (206) 2024 | x | - | - | x | - | x | 2 | 2 | 3 | active |

## Results

Our systematic review highlights the diversity of multi-omics literature in terms of research focus and methodology. We categorized the reviewed studies into “host and host-associated microbiome”, “microbiome-free host” and “host-free microbiome” data (Fig. 3; Table 1, Table 2, Table 3). These summarize the distribution of categories, sample types, data types, host species and analysis methods. Overall, the types of multi-omics data and associated computational methods in these studies ranged from general exploratory techniques to more advanced time-series-specific methods designed for longitudinal datasets. Common study designs for longitudinal studies included monitoring studies, cohort studies and intervention studies. Cohort studies track a sample or cohort of randomly selected individuals from a homogeneous group over time, frequently documenting the progression of the disease or natural variability. These studies are useful for finding patterns or biomarkers linked to certain outcomes, including the start or recovery from illness (16; 17; 18). Monitoring studies involve the ongoing or sporadic monitoring of participants for an extended period of time, frequently in uncontrolled or natural settings. Understanding the impact of changes in the environment or lifestyle variables is made easier by such studies (19). Intervention studies compare the time-series data across two or more groups undergoing different treatments, such as clinical trials or dietary interventions. These studies are especially helpful for determining how certain therapies affect multi-omics profiles over time (20; 21; 22; 23; 18; 24).

**Fig. 3.**
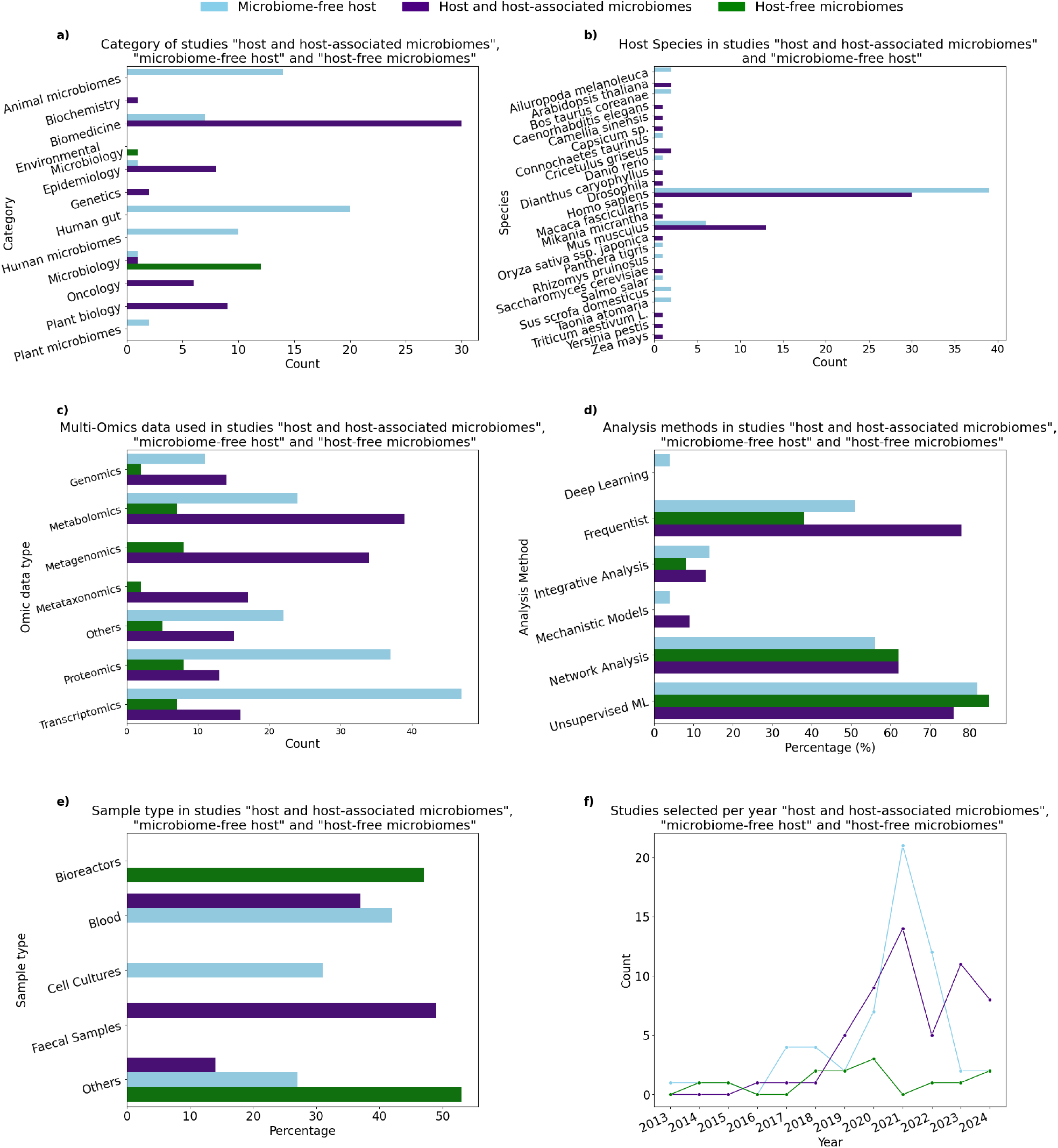
Comparative analysis of studies: “host and host-associated microbiome”, “microbiome-free host” and host-free microbiome”: (a) distribution of study categories, (b) number of host species identified, (c) types of ’omics data used, (d) Percentage of analytical methods applied, (e) sample types analysed, (f) number of studies published from 2013 to 2024.

**Fig. 4.**
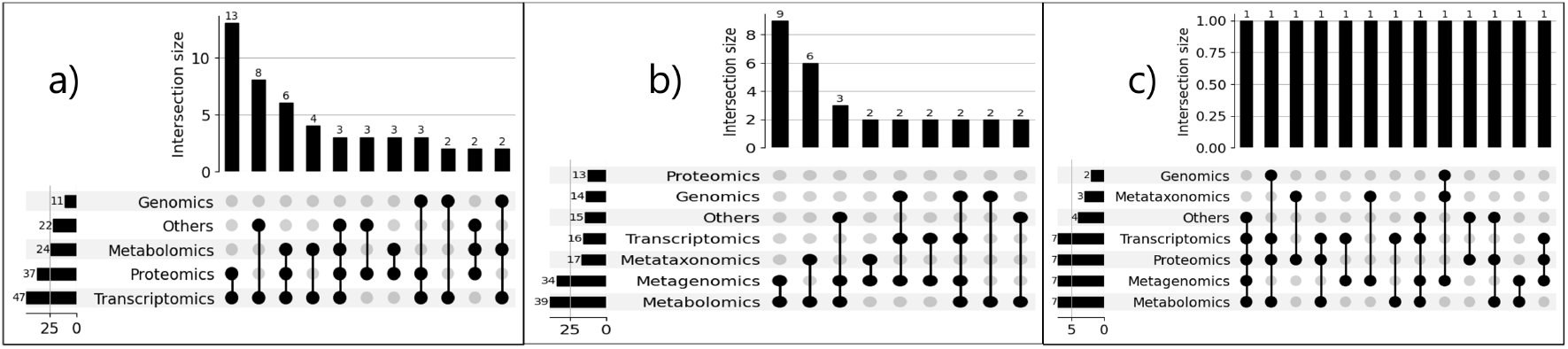
Upset plot visualizing the overlap of multi-omics data type across the reviewed studies: (a) “microbiome-free host” with intersection size = 2, (b) “host and host-associated microbiome” with intersection size = 2 and (c) “host-free microbiome” with intersection size = 1.

## Methodological studies

We identified in total 18 studies which described modeling frameworks for multi-omics time-series analysis (Table 4). The most common analysis method categories included in the 15 frameworks are ordination or unsupervised machine learning (8 studies), frequentist uni/multivariate methods (7 studies) and network analysis (7 studies).

We evaluated the methods based on three key aspects: performance, interpretability and ease of use. These aspects were qualitatively evaluated with scores from 1 (worst) to 3 (best). Based on this qualitative assessment, we noticed shortcomings in one or more of these criteria in two studies, whose repositories have been deprecated or did not received updates since 2014. The methods developed by Chong et al. (25), Conard et al. (26) and Hamzeiy et al. (27) achieved maximum scores; all of these methods are still actively maintained. The methodology developed by Chong et al. (25) showed the broadest application in multiomics time-series analysis (28; 29; 30; 31; 32). Other methods were frequently cited, but their usage and applications were not clearly described. This indicates a potential gap in reporting or less-defined roles in practical analyses. Interestingly, some of the methods appeared to have been available for a long time before the study itself was published. For example, the code repository of ADAPT (Analysis of Dynamic Adaptations in Parameter Trajectories; van Riel et al. (33)) was last updated in 2014, while the study was published in 2021.

### Multi-omics data

Although including multiple ’omics layers offers advantages in microbiome research, the consensus on the optimal approach to achieve the integration of different ’omics layers in the inference framework is yet to be achieved. Some studies use many different aspects of microbiome observations, such as metagenomic, meta-transcriptomic and metaproteomic data, capturing different aspects of the biological processes (e.g., taxonomic, potential and realized function). In some cases, data pre-processing involved performing batch correction by combining information across these multiple ’omics layers. In other cases, data from many ’omics were combined for downstream analysis. Traditionally and most simply, several different studies have been performed, so that each ’omics layer is analysed independently, followed by qualitative discussion about the observed parallel changes.

### Dimensionality reduction

Dimensionality reduction is commonly used as the initial stage of most of the multi-omics studies, because it enables exploratory analysis and visualization of the dataset. Principal Component Analysis (PCA) a widely used method for dimensionality reduction, is frequently applied to individual ’omics datasets to extract features or reduce noise prior to integration (34; 35; 36; 37; 38; 39; 40; 41; 42; 43; 44; 45; 46; 47; 48; 49; 50; 51; 52; 23; 53; 54; 55; 56; 57; 58; 59; 60; 61; 62; 20; 63; 64; 65; 66; 67; 68; 69; 22; 70; 71; 72; 73). For basic integration, some methods employ PCA immediately after concatenating abundance tables from many ’omics (such as transcriptomics and proteomics) into a single matrix. Pipelines have been shown to use PCA to find latent features for downstream classification tasks using combined metagenomic and metabolomic data (74).

Principal coordinate analysis (PCoA) is essentially a form of classical multidimensional scaling (MDS) that extends PCA to non-Euclidean dissimilarity measures and a common choice in microbiome research. This was the second most used method in the studies included in this review (46; 75; 76; 42; 77; 51; 78; 79; 53; 80; 81). In time-series multi-omics, these methodologies enable temporal trajectory studies by condensing data variance between time points into a reduced number of interpretable dimensions. MDS has also been employed in multi-omics integration by computing joint dissimilarity metrics across ’omics layers, though this may require careful normalization to balance feature scales.

Furthermore, alternative (nonlinear) dimensionality reduction methods have also been used, including Isomap, t-SNE (45; 82; 83; 84) and UMAP (85; 62; 83; 86; 22; 87). While these are often applied to single-omics data, recent workflows integrate multi-omics by first reducing each layer separately using PCA and then aligning embeddings. For example, Compound-SNE aligns t-SNE projections from multiple single-cell ’omics datasets while preserving sample-specific structures (88). Such methods address the limitations of naive concatenation by leveraging shared variance or feature-grouping strategies. In summary, while PCA, MDS or t-SNE are frequently applied to individual ’omics layers, we also identified their use in multi-omics integration, either through concatenation or coordinated embeddings.

### Correlation analyses

The choice of the methods is often based on specific data characteristics, including the type of data (e.g. continuous or categorical), data distribution and measurement scale. For example, pairwise correlation coefficients have been calculated between relative abundances of microorganisms and the expression levels of several genes. Several studies used Pearson’s correlation (89; 24; 47; 90; 91; 57; 92; 93; 94; 68; 95; 84). In longitudinal multi-omics, such correlations can be used to monitor the evolving associations between molecular variables across time, aiding in the identification of consistent or temporary interactions across temporal points. However, other studies utilized Spearman correlation (49; 96; 97; 98; 99; 100; 81; 101; 68; 102; 72) which involves transforming features into ranks and can better quantify non-linear associations. Rank-based methods such as Spearman’s correlation can thus be very useful as they can identify monotonic trends over time.

The possibility of compositionality bias must be taken into account when determining correlations for compositional data, such as relative abundances of microbiomes. By definition, compositional data add up to a constant (100% relative abundance), therefore modifications to one component always impact the others. Pearson and Spearman correlations are subject to this bias, unless specifically corrected for it(103). The centered log ratio (CLR) transformation is often used to mitigate compositional effects by converting compositional data into a log ratio space. The CLR transformation calculates the logarithm of the abundance of each trait relative to the geometric mean of all traits in the sample in order to mitigate dependencies between features (104). Whereas the use of CLR or comparable transformations is increasingly recognized as standard practice in microbiome research to ensure reliable correlation analyses, this issue was not always addressed in the reviewed studies.

Furthermore, canonical correlation analysis, an extension of the principal component analysis to multiple datasets (105; 106), can quantify multivariate correlations between datasets. It can identify correlated feature sets in paired datasets, instead of individual correlated pairs of individual features detected by the standard Pearson and Spearman cross-correlation. This approach has been recently used, for example, by Revilla et al. (107) and Simats et al. (60).

### Clustering & similarity network methods

Clustering methods are used to discern general patterns in a dataset. Clustering can be performed on both samples and features. Some of the reviewed studies used dissimilarity measures such as Euclidean distance, Manhattan distance or Bray-Curtis dissimilarity (38; 39; 79; 18; 80) to perform clustering across samples. Clustering techniques can be used for basic integration by defining distance metrics across several multi-omics layers. Integrative techniques that are relevant to multi-omics time-series analysis include iCluster (108) and Similarity Network Fusion (SNF) (109). These methods are designed to handle the complexity of multi-omics data by capturing both shared and layer-specific patterns across time.

SNF is a powerful integrative method that constructs and fuses sample similarity networks across multiple ’omics layers. SNF functions in the sample space, as opposed to feature-based networks, where nodes indicate samples (such as patients or time points) and edges indicate pairwise similarities between samples according to their ’omics profiles. To create a unified representation that captures both shared temporal patterns (common across ’omics layers) and layer-specific temporal patterns (unique to a particular ’omics layer), SNF builds distinct similarity networks for each ’omics layer (e.g., transcriptomics, proteomics) at each time point. This fusion approach is especially useful for detecting dynamic biological changes that are consistent across various data types because it makes use of local commonalities and complementary information across ’omics layers. In cancer research, Wang et al. (110) used SNF to integrate data on mRNA expression, DNA methylation and miRNA expression. This has revealed temporal trajectories and clinically significant subgroups that were not visible in individual ’omics layers.

In contrast, iCluster concentrates on separating the data into shared and distinct patterns that show temporal dynamics, such as metabolites that only show up at particular times, or gene expression levels that fluctuate over time. iCluster thus finds sample clusters that change over time in response to biological disturbances, such as the course of a disease or the results of therapy, by modelling these time-specific properties. Shen et al. (111) utilized iCluster as a joint latent variable model that combines transcriptomic, proteomic, epigenomic and genomic data to categorise tumor subtypes. iCluster outperformed conventional separate clustering techniques.

### Regression & classification

Regression and classification techniques can be used for asymmetric quantification of associations between two ’omics, for instance to predict values of one set based on the other. Many studies have focused on the case of a single ’omics regression (i.e., predicting one ’omics layer as the response from another as the predictor, *y ~ x*). However, an equally important and emerging direction involves integrating two or more ’omics layers to predict external covariates such as age, BMI, or overall health. In the simplest approach, datasets from different ’omics layers can be concatenated. Several studies used linear mixed models (112; 37; 38; 76; 44; 18; 53; 113; 82; 99; 114) to establish associations between ’omics layers. Several studies have explicitly applied multi-omics regression and classification approaches in time-series settings. For instance, studies integrating metagenomic and metabolomic datasets have concatenated data to predict host phenotypes over time, thereby revealing dynamic associations that evolve with aging or health status (46; 90).

For classification, several studies used Linear Discriminant Analysis (LDA) (43; 77; 96; 113; 59). This method identifies the most optimal hyperplane to separate labeled samples. Also, extended discriminant analysis method called sparse variant (sPLS-DA) was used in several studies (39; 115; 48; 49; 52; 79; 98; 116; 72). sPLS-DA performs variable selection and classification in a one-step procedure and enables the selection of the most predictive or discriminative features in the data to classify the samples. Several studies utilized individual ’omics even within multi-omics studies. However, recent studies have extended these classification approaches to directly integrate multiple ’omics layers. In such frameworks, sPLS-DA has been successfully used to track time-evolving discriminative features across transcriptomic, proteomic and metabolomic data, thus enhancing the predictive and interpretative power in longitudinal studies (72; 79). Linear mixed models efficiently incorporate random effects related to temporal variability, thereby enabling researchers to rigorously evaluate longitudinal trends in multiomics associations.

### Temporal modeling & longitudinal data analysis

Various methods have been specifically devised for longitudinal multi-omics. One of the main challenges in multi-omics integration is handling asynchronous sample intervals and disparate progression rates across various ’omics layers. Dynamic Bayesian Networks (DBNs) are especially useful in this context, as they determine directed connections among biological entities - such as host genes, metabolites and microbial taxa - while capturing the non-linear and conditional dependencies present in biological systems. Vector autoregressive models presume linearity and although recurrent neural networks (RNNs) can achieve high prediction accuracy, they frequently operate as “black boxes” that lack interpretability. Ruiz et al. (117) addressed these challenges by proposing the PALM pipeline. This approach first aligns longitudinal data from host transcriptomics, metabolomics and metagenomics and then uses DBNs to reconstruct a unified interaction network. The PALM pipeline has effectively identified both known and novel metabolite–taxon interactions in patients with inflammatory bowel disease (IBD), with experimental validation further supporting these findings.

Other approaches to modeling temporal dependencies across ’omics layers include state-space models (118) and vector autoregressive models (119). Certain variants of RNNs, such as long short-term memory (LSTM) networks, are capable of capturing temporal patterns in multi-omics data (120). Additionally, network-based methods—including temporal correlation networks and multilayer networks—can combine and examine patterns in multi-omics time-series, emphasizing the dynamical transitions and trends (121).

### Neural networks & deep learning

The integration of heterogeneous multi-omics data collected over time has been enabled by recent advances in deep learning, providing unprecedented insights into biological systems and disease processes. Jain and Safo (122) developed a deep learning pipeline that uses gated recurrent units (GRUs) to extract time-dependent features for disease classification, thereby integrating cross-sectional and longitudinal multi-omics data, including transcriptomics, metabolomics and metagenomics. It stands out for its ability to handle non-overlapping samples and variable-length time-series data, maximizing the use of available heterogeneous datasets.

The convolutional graph neural network (ConvGNN) framework for multi-omics categorization of chronic obstructive pulmonary disease (COPD) was established by Zhuang et al. (123) as a complementary method. Unlike traditional classifiers, this study improves prediction accuracy by combining protein-protein interaction (PPI) networks from known databases with longitudinal proteomic and transcriptomic data. The ConvGNN technique improves the interpretability and efficiency of COPD classification models by integrating biological network information into the learning process.

Furthermore, Lim and van der Schaar (124) introduced Disease-Atlas, a deep learning technique that simultaneously models time-to-event outcomes and longitudinal data. This method enables more accurate predictions of disease progression by using adaptive neural network architectures to capture the dynamic evolution of disease states from multi-omics inputs.

### Multi-omics (latent) factor analysis

Multi-omics factor analysis (MOFA) is a powerful framework designed to separate variation in complex multi-omics datasets by providing a shared low-dimensional representation that captures common and modality-specific signals. Argelaguet et al. (125) used cross-sectional cohort of chronic lymphocytic leukaemia (CLL) patient samples, where MOFA integrated somatic mutations, RNA expression, DNA methylation and ex vivo drug responses to uncover major dimensions of disease heterogeneity (such as immunoglobulin heavy-chain variable region status and trisomy of chromosome). MOFA has proven invaluable for revealing underlying biological processes in complex multi-omics datasets.

Several studies have extended MOFA to address the difficulties presented by longitudinal data based on this foundation. Zimmer et al. (45) analysed longitudinal multi-omics data including proteomics, metabolomics, microbiome and clinical laboratory values, using the Pareto Task Inference (ParTI) approach. This method showed that three wellness stages and one aberrant health condition were defined by the mapping of clinical lab data onto a tetrahedral structure. Similarly, MOFA was used by De et al. (126) on a longitudinal murine model. Their analysis revealed that gut microbial and metabolic alterations, particularly in bile acid, energy and tryptophan metabolism, preceded allergic inflammation following *β*-lactoglobulin (BLG) sensitization. These findings were validated in children with IgE-mediated cow’s milk allergy (IgE-CMA), linking gut dysbiosis to early immune responses. This highlights microbial and metabolic markers as potential early predictors of IgE-CMA.

Gaussian process regression is integrated into MEFISTO to model spatio-temporal dependencies in longitudinal multi-omics data, extending traditional factor analysis frameworks. In their foundational work, Velten et al. (127) applied MEFISTO to evolutionary developmental atlases (gene expression data from five species across organ development), longitudinal microbiome studies (43 children over two years) and single-cell multi-omics datasets (mouse gastrulation with RNA, methylation and chromatin accessibility). These applications revealed conserved developmental trajectories, species-specific variation and dynamic gene regulation, outperforming conventional methods in imputing missing data and aligning temporal patterns across misaligned groups. MOFA+, the framework underpinning MEFISTO, extends these capabilities to integrate multimodal single-cell data across diverse sample groups. MOFA+ has been used to model heterogeneity in immune-mediated diseases by jointly analyzing DNA methylation, chromatin accessibility and transcriptomic profiles, identifying latent factors linked to dynamic T cell activation states (128). This approach leverages computationally efficient variational inference to unify large-scale and single-cell datasets, enhancing patient stratification through temporal or disease progression-associated features.

## Discussion

### Time-series data collection

Microbiomes are inherently dynamic; therefore, gathering and analysing longitudinal data is necessary to better understand the interactions within host-microbiome communities. Such studies can help us to better understand complex mechanisms between the multi-omics profile of an organism and its phenotype, as well as how biological systems respond to variations in their genetic makeup or external environments. Including multiple samples in the analysis allows us to identify the essential core interactions between a host and its microbiome. This approach also provides a unique opportunity to quantify the correlation or divergence between time points and compare these metrics across the different layers of ’omics.

Collecting time-series data in the context of multiomics poses distinct and considerable challenges. Obtaining consistent sampling across these domains at regular intervals is especially challenging when investigations span extended periods. Moreover, regulating environmental and experimental variability is difficult due to the dynamic nature of living systems. Gene expression, protein synthesis and metabolic activity can change unpredictably, even under steady conditions, causing variability that can hinder data interpretation. Researchers are thus increasingly implementing stringent criteria for sample handling, storage and archiving to ensure uniformity over time and between study sites (129). Ethical and logistical constraints introduce additional complications, particularly in studies involving humans or animals. Repeated sampling may be impractical due to ethical considerations or the intrusive nature of the methods. To overcome these obstacles, researchers frequently employ “pseudo time-series” approaches by sampling distinct individuals (yet if possible similar in the major characteristics) at various time intervals and merging the data to deduce temporal trends (130; 131). While this can provide useful insights, it cannot match the depth of information obtained by monitoring changes within the same individual over time. Consequently, it may overlook nuanced biological rhythms or fail to adequately document the comprehensive development of diseases. Designing an efficient time-series study requires achieving a careful balance among sampling frequency, temporal resolution and the ethical constraints associated with the study’s subjects and aims.

By integrating host and microbiome data across multiple time points into a unified framework, we can maximize its potential. This approach enables our understanding and accurate prediction of dynamic phenotypic traits, including growth dynamics, health, drug response, disease susceptibility and pathogenesis (119).

### Data types and data structures

Multi-omics data is often sparse due to many practical and ethical challenges related to experimental design and sample collection. A significant issue is the lack of one-to-one matching across different ’omics layers, meaning that not all samples are measured across all modalities (e.g., genomics, transcriptomics, proteomics). This results in unevenly distributed and missing data, which can limit the robustness of conclusions drawn from such datasets. A typical multi-omics study might examine six major technique categories: genomics, transcriptomics, proteomics, metabolomics, epigenomics and single-cell ’omics. However, due to technical limitations, cost, or sample availability, only a subset of these techniques is often applied, leading to incomplete data integration and potential biases in analysis. Hence, it weakens the opportunity of comparing different studies since they collect different type of data.

The arrangement of data in appropriate containers and formats plays a significant role in managing multiomics time-series datasets. Efficient data storage and retrieval technologies provide tools to readily access, process and analyse data across different ’omics layers. Contemporary data storage formats, such as HDF5, OME-Zarr (132), OME-NGFF (133) have emerged as favored choices due to their capacity to manage extensive, multi-dimensional datasets effectively (134). The R/Bioconductor community has advanced statistical data analysis methods based on specific multi-assay data structures (135; 136). These and other formats support multi-source data integration and can facilitate hierarchical data organization, permitting researchers to consolidate many types of ’omics data within a singular container while preserving their unique structures and formats. Moreover, multi-omics data retrieval tools (e.g. HoloFoodR (137)) and interactive applications (e.g. iSEEtree (138)) support the exploration and analysis of longitudinal and other multi-omics datasets based on such data structures.

Interoperability across diverse data formats and platforms is especially crucial in multi-omics research, as it enables for smooth integration of datasets from different sources or studies. Standardized formats like JSON and XML for metadata annotation assist in maintaining compatibility, enabling researchers to correlate data on gene expression, protein levels, metabolite concentrations and other factors across time points. This interoperability is crucial in collaborative studies with multi-site or multi-disciplinary teams that contribute data to a common repository. The utilization of modular and adaptive data containers facilitates data accessibility, retention and reproducibility, thus facilitating deeper insights into host-microbiome interactions over time.

### Underutilized analysis techniques

Mechanistic modeling of multi-omics measurements holds the promise of providing a more comprehensive and nuanced representation of biological systems, when compared to data-driven inference and deep learning methods. Mechanistic models, such as dynamic models employing differential equations, or agent-based models, could encapsulate key aspects of a system’s behavior (139). Such approaches have been previously used to elucidate molecular interactions, gene regulatory networks and causal linkages (140). Moreover, they have demonstrated utility in uncovering regulatory mechanisms in both healthy and pathological conditions, as well as in examining recovery processes from disrupted states (26). Thus, mechanistic models can help establish a solid basis for refining interactions, assessing and validating ranges of kinetic parameters, identifying most important model components and to better understand the underlying mechanisms and drivers of microbiome dynamics.

The computational strategies for integrating longitudinal multi-omics data are only starting to emerge. As previously highlighted, separate analysis and post-hoc comparisons of multi-omics data are often inadequate for gaining deeper insights into the interactions between the different ’omics layers. Integrative techniques are essential for understanding interactions across diverse biological processes and ’omics data. The intrinsic variability and irregular data availability multi-omics time-series underscore the necessity for adaptability in analytical frameworks. Whereas traditional methods often presume comprehensive and uniformly distributed data, biological data often display deficiencies or uneven temporal intervals due to logistic limitations or sample attrition (141). Adaptive techniques that can tackle these issues are crucial for producing significant discoveries. Methods like imputation of absent values, interpolation models, Bayesian techniques and machine learning algorithms have arisen as essential instruments in this domain. Machine learning techniques, including recurrent neural networks (RNNs) (142; 143), long short-term memory (LSTM) models (144) and transformers (145; 146), show great potential for modeling temporal correlations in multi-omics data, even when faced with missing or irregularly spaced observations. While transformers have not yet been extensively applied to time-series data (to the best of our knowledge), their inherent memory mechanisms and ability to capture long-range dependencies suggest they could be highly effective for modeling such data in the future. Nonetheless, these models frequently need substantial computational resources and specialised knowledge, which may restrict their wider utilisation. Furthermore, the amalgamation of diverse data types in time-series analysis continues to provide a significant difficulty. Each ’omics data type displays unique properties to consider. Thus, enhancing the adaptability of analytical tools will be essential for realising the complete potential of longitudinal multiomics. As these databases grow in complexity and scale, adaptive methods will be essential for enabling comprehensive analyses.

### Adoption gap

Despite the availability of the proposed methods, their integration into widely utilized computational frameworks remains limited. In many cases, either the implementations are unavailable, or they are restricted to specific software environments that may not be accessible to all researchers. Furthermore, many methods are tailored to particular use-cases, making them challenging to adapt for other types or collections of data. This has resulted in an “adoption gap” of the new methods. Cross-disciplinary training programs could support the broader computational application and development of skills among applied researchers. The creation of intuitive graphical user interfaces and streamlined workflows in widely used platforms and cloud-based technologies might further enhance the adoption of these methods.

### Challenges and limitations in multi-omics data integration

The interactions between the biological processes of the host and their microbiome are still only superficially understood. Integrative analysis of the (meta) genomes, (meta) transcriptomes and (meta) metabolomes of the host and its microbiomes is a more extensive approach than analyzing each of these ’omics data separately (147). Creating a comprehensive framework that combines data from many ’omics layers and time intervals enables more effective discovery of biological pathways that link, e.g., genomic variation to phenotypic variance. By cross-comparing different ’omics layers, we can examine direct interactions within these layers (*e.g.*, host genome to metabolome) and between the host and its microbiomes. This allows us to comprehensively and systematically understand the intricate biology that underlies the connections between the host genome and health, as well as the composition or diversity of the microbiome (148).

Several limitations and biases in the reviewed studies and their methodologies remain despite the potential of multi-omics integration. Some of the main challenges include the relatively small number of time points, which may be further unevenly spaced or unmatched between different data types, high individual variability and subject drop-outs (149). Sometimes it is not possible to collect longitudinal data, for example, because the sample is drawn from tissue or an organ which is surgically removed. Moreover, invasive sample collection at more than one time point might not be ethically justified and there are high costs involved with sampling at multiple time points. In the case of laboratory animals, it is possible to collect samples that require euthanizing; however, this design does not allow samples from multiple time points to be collected. In these cases, a so-called “pseudo time-series” can be assembled from multiple cross-sectional datasets so that, for example, disease progression is preserved (see, e.g., (15)). This means that at each time point, the disease state is carefully identified and the full dataset consists of ordered time points which simulate disease progression. Pseudo time-series can thus approximate the collection of true time-series data in these cases. However, intra-individual differences might disguise patterns related to disease progression.

A large number of studies include only a few entities that were tracked over time, especially in the context of “host and host-associated microbiomes”. For example, the two studies on swine (*Sus scrofa domesticus*) microbiomes sampled only three (150), or six (151) animals over time. Furthermore, the main problem is generally not limitations in sample size per se but the level of heterogeneity.

### Implications for future research

In summary, an interdisciplinary data integration strategy should be used to support a better understanding of hierarchically structured complex biological systems. This would enable predicting trajectories of change, optimizing the predictive power of theoretical models and developing successful practices for agriculture, aquaculture, veterinary science and human health. A better understanding of genotype-phenotype associations, as well as the biological pathways between them, will allow us to identify better interventional targets in biological systems, such as better probiotics in food production systems within agriculture and aquaculture or gene targets for drugs. It will also allow us to develop precision medicine and predict future changes in the microbiome in response to such treatments (152; 153).

## Perspectives

Accounting for the temporal dimension in multiomics studies is a rapidly expanding research theme.

However, the heterogeneity of analytical approaches in current studies and the need for more systematic approaches centering around specific well-defined application tasks is clear. There is a rapidly increasing need for integrative analysis methods and open research software. Such tools are essential for supporting the practical application of the many rigorous statistical and machine learning methods recently introduced in this research area. Results from such studies could be expected to have an increasing impact in ecological, evolutionary and medical research.

## Code and data availability

All the code and data tables used for the figures in the manuscript are available on github here.

## Key Points

- Time-series multi-omics studies are becoming the standard for studying temporal and functional aspects of host-microbiome systems.
- Most studies use only exploratory analyses for summarizing time-series multi-omics data.
- Only a few integrative frameworks exist for analysing time-series multi-omics data.
- This study presents an overview of the current methods and techniques, thus providing a pipeline for time-series studies starting from data collection to integrative inferences.

## Acknowledgments

This project has received funding from the European Union’s Horizon 2020 research and innovation programme under grant agreement No 952914.

## Author contributions statement

M.K.S., M.O.R., S.G. and L.L. conceived the study. M.K.S. and M.O.R. collected the data, while all authors annotated the collected articles with relevant information. M.K.. and M.O.R. drafted the manuscript, and S.G. and L.L. provided extensive critical revisions and oversaw the manuscript preparation throughout the writing process. Selected studies were divided to D.F.B., P.N.F., G.B., A.M., T.B., P.P.E and I.S. to cross-check for relevance and accuracy. All authors contributed to data interpretation, commented on the manuscript, and approved the final version for submission.

**Moiz Khan Sherwani** received his Ph.D. in Computer Science with a specialization in AI in medical imaging across diverse modalities. He is currently a postdoctoral researcher at the University of Copenhagen, where his research focuses on developing advanced deep learning techniques including Graph Neural Networks (GNNs) and Convolutional Neural Networks (CNNs) for the analysis of multi-omics data, as well as applying rigorous statistical methodologies to derive meaningful biological insights. **Matti Ruuskanen, PhD,** has studied microbial communities in both environmental and host-associated settings. He has published papers on lake sediment microbiomes, human gut microbiome and its associations with chronic diseases, and microbiome data science. He is currently employed as a University Lecturer at the Department of Life Technologies at the University of Turku.

**Dylan Feldner-Busztin** studied finance as an undergrad before making the switch to neuroscience as a masters. Since then he has been working in various domains of computation biology, including machine learning for bioinformatics and agent-based modelling for cell migration.

**Panos Fibras** studied Biology and computational science before doing a PhD on the evolution of cis-regulation through bioinformatics and machine learning. He focuses on working with a large variety of -omic data and applying statistical and ML techniques to gain biological insights.

**Gergely Boza** is a theoretical biologist working at the Institute of Evolution in Hungary, and at the International Institute for Applied Systems Analysis (IIASA) in Austria. He is in general interested in modeling complex systems, from microbial ecosystems to socio-economic systems, including the evolution of cooperation and mutualisms. Currently his main research focuses on developing individual-based models of microbiomes.

**Ágnes Móréh** is a theoretical ecologist specializing in the dynamic modeling of ecosystems. Her research focuses on the network aspects of biological invasions and the mathematical modeling of superinfections. She employs mathematical and computational modeling techniques to analyze the complex interactions within ecological systems.

**Tuomas Borman** is a doctoral researcher at the University of Turku, working as part of the Turku Data Science Group. His research focuses on developing microbiome data science methods within Bioconductor.

**Pande Putu Erawijantari** is a postdoctoral researcher specializing in microbiome and multiomics research. Her research focus explores the role of gut microbiomes in health and disease, from treatment outcomes to population-wide health trends.

**István Scheuring** is a theoretical biologist interested in evolutionary ecology, mutualism, and microbiome dynamics in general. Within microbiome ecology, nowadays he focuses on modeling microbial communication, cooperation, and host-microbiome interactions.

**Shyam Gopalakrishnan** is an associate professor at the Globe Institute, University of Copenhagen. His research focus is multi-omics integration, evolutionary and population genomics. He focuses on developing methods for integrating data from different biological processes in host-microbiome systems, with special interest in modeling co-evolution of microbiomes together with their hosts.

**Leo Lahti** is professor in Data Science at the Department of Computing, University of Turku. His main research focus is on computational microbiome research. The research team has developed algorithmic methods and Bioconductor data science frameworks for multi-omic data integration in this research area.

## Notes

https://github.com/shyamsg/TimeSeries_MultiOmics_Review

### Competing Interest Statement

The authors have declared no competing interest.

